# Highly accurate isoform identification for the human transcriptome

**DOI:** 10.1101/2022.06.08.495354

**Authors:** Markus J. Sommer, Sooyoung Cha, Ales Varabyou, Natalia Rincon, Sukhwan Park, Ilia Minkin, Mihaela Pertea, Martin Steinegger, Steven L. Salzberg

**Affiliations:** Department of Biomedical Engineering, Johns Hopkins School of Medicine and Whiting School of Engineering; Baltimore, MD, United States; Center for Computational Biology, Johns Hopkins University; Baltimore, MD, United States; School of Biological Sciences, Seoul National University; Seoul, Korea; Artificial Intelligence Institute, Seoul National University; Seoul, Korea; Department of Computer Science, Johns Hopkins University; Baltimore, MD, United States; Institute of Molecular Biology and Genetics, Seoul National University; Seoul, Korea; Department of Biostatistics, Johns Hopkins University; Baltimore, MD, United States

## Abstract

We explore a new hypothesis in genome annotation, namely whether computationally predicted protein structures can help to identify which of multiple possible gene isoforms represents a functional protein product. Guided by structure predictions, we evaluated over 140,000 isoforms of human protein-coding genes assembled from over 10,000 RNA sequencing experiments across many human tissues. We illustrate our new method with examples where structure provides a guide to function in combination with expression and evolutionary evidence. Additionally, we provide the complete set of structures as a resource to better understand the function of human genes and their isoforms. These results demonstrate the promise of protein structure prediction as a genome annotation tool, allowing us to refine even the most highly-curated catalog of human proteins.

**One-Sentence Summary:** We describe the use of 3D protein structures on a genome-wide scale to evaluate human protein isoforms for biological functionality.

## Main Text

More than twenty years after the initial publication of the human genome, the scientific community is still trying to determine the complete set of human protein-coding genes. Although the number of genes is converging around 20,000, we do not yet have agreement on the precise number. The true number of different isoforms of human genes – variations due to alternative splicing, alternative transcription initiation sites, and alternative transcription termination sites – is even less certain. Currently, the major human gene annotation databases each contain well over 100,000 protein-coding isoforms (*1–5*), but the sets of isoforms vary widely among them.

Although the functions of many human genes are known, elucidating gene function remains a complex and time-consuming task. Given that at least 92% of human genes express more than one isoform (*6*), and that the human transcriptome contains an average of seven or more unique transcripts per protein-coding gene (*7*), the only feasible way to determine which isoforms are functional on a genome-wide scale is by using computational methods. Until now, the primary computational tools used to investigate gene function were sequence alignment and gene expression. Alignment relies on the long-established observation that if a protein is conserved in other species, then it is likely to be functional, particularly if the conservation extends to distantly-related species (*8*). This rule applies to isoforms as well: if we can find evidence that a particular sequence – e.g., a protein that uses an alternative exon – is present in species that diverged tens of millions of years ago, then the conservation of the sequence argues in favor of its function.

In a similar vein, the use of RNA sequencing (RNA-seq) to detect gene expression also provides clues to function: if a transcript is consistently expressed in multiple samples, it is more likely to be functional than one for which no expression evidence can be found. Genes may encode multiple transcripts that fold into distinct isoforms with well-defined functions (*6*), but recent work has shown that most assembled human transcripts are found at very low levels in the transcriptomes of individual tissues (*4*), and may simply reflect biological noise, products of intrinsically stochastic biochemical reactions (*9, 10*). The large majority of assembled isoforms are unlikely to be functional, and indeed only a small percentage are included in current human genome annotation databases (*1–5*). Because transcription is noisy, the observation of transcription in RNA-seq data is insufficient evidence to conclude that a sequence is functional (*11*).

This study explores a fundamentally new line of evidence that can be used to investigate protein function: computational prediction of three-dimensional structure. The recently-developed AlphaFold2 system can automatically predict three-dimensional protein structure with accuracy that often matches far more time-consuming laboratory methods (*12, 13*), allowing us to generate structure predictions for thousands of gene isoforms. In proteins where a substantial portion folds into an ordered structure, estimated to be 68% of human proteins (*13, 14*), a well-folded structure within an isoform argues in favor of its functionality. Conversely, a poorly-folded isoform may indicate loss of function.

In a recent effort to create a single consensus annotation of all human protein-coding genes, two of the leading human genome annotation centers created the MANE (Matched Annotation from NCBI and EMBL-EBI) database (*15*), a high-quality collection of protein-coding isoforms for which the annotation databases RefSeq (NCBI) and Ensembl-GENCODE (EMBL) match precisely. The goal of MANE is to identify just one isoform for each protein-coding gene that is well-supported by experimental data, and to ensure that both databases agree on all exon boundaries as well as the sequence of the associated protein. In addition to the one-isoform-per-gene collection known as MANE Select, a small number of additional transcripts with special clinical significance, known as MANE Plus Clinical, are included in the database. Upon its initial release, MANE included only around 50% of human protein-coding genes. The latest version, v1.0, includes 19,062 genes and 19,120 transcripts, with an additional 58 transcripts included in the MANE Plus Clinical set. These transcripts have been described as a “universal standard” for human gene annotation, and they provide a valuable resource to scientists and clinicians who need a consistent set of functional primary transcripts.

Here we describe our use of protein folding predictions from ColabFold (*16*), an open-source accelerated version of AlphaFold2, alongside experimental RNA-seq expression data from the Genotype-Tissue Expression project (GTEx) (*17*), to present substantial evidence for functional isoforms that can be used to improve human gene annotation, including the MANE gene set as well as the comprehensive human annotation databases RefSeq (*2*), GENCODE (*3*), and CHESS (*4*). We dive into a few exemplary predictions to explain, biologically and evolutionarily, the three-dimensional structure of our alternate isoforms. We also present an example of a novel protein isoform in mouse to demonstrate the general applicability of this structure-guided approach to improving functional annotation of any genome.

## Results

### Scoring the transcriptome

Using a large set of transcripts assembled from nearly 10,000 human RNA-sequencing experiments, we identified all isoforms of protein-coding genes that were 500aa or less in length (see Methods). The 142,280 transcripts at 20,021 gene loci that fit this description encoded 90,413 distinct protein isoforms, and we predicted structures for all of them. Gene identifiers for all predicted protein isoforms as well as predicted Local Distance Difference Test (pLDDT) scores and evolutionary conservation data from mouse can be found in Table S1. Predicted scores and GTEx expression data for all isoforms overlapping a MANE locus can be found in Table S2. All predicted protein structures as well as data for all tables in this paper are publicly available at our website, isoform.io.

After comparing all predicted structures to those for proteins in MANE, we identified 401 MANE loci that contained at least one alternate isoform that appeared to have a more stable structure than the annotated primary isoform. Data for these 401 alternate isoforms can be found in Table S3. Worth noting is that over 95% of the MANE loci evaluated here contained no higher-scoring alternate isoforms that passed our filtering criteria. This is a testament to both the high degree of consistency in MANE as well as the sensitivity of protein structure prediction for finding instances where alternative isoforms create functional products.

### Exemplary predictions

To illustrate the improvements in human gene annotation that can be obtained using accurate structure prediction, we describe a small set of proteins, selected from Table S3, where an alternate isoform appears to be superior to the isoform chosen for inclusion in MANE. For these examples, the alternative isoform is clearly functional based on structure as well as evolutionary conservation and, in some cases, additional expression evidence from RNA-sequencing data. For some of these examples, the MANE isoform is missing critical structural elements and may not be functional at all.

#### Acetylserotonin O-Methyltransferase

Acetylserotonin O-methyltransferase (*ASMT*, alternatively *HIOMT*) is responsible for the final catalytic step in the production of melatonin, a critical hormone in sleep, metabolism, immune response, and neuronal development (*18*). Depressed levels of circulating melatonin have been associated with autism spectrum disorder, and clinical studies have classified *ASMT* as a susceptibility gene due to the highly significant association between *ASMT* activity and autism (*18, 19*).

The CHESS and GENCODE gene databases contain a 345aa isoform of *ASMT* (CHS.57426.4, ENST00000381229.9) while RefSeq is missing this isoform. The MANE version of this gene is 373aa long and appears in the CHESS (CHS.57426.2), GENCODE (ENST00000381241.9), and RefSeq (NM_001171038.2) gene databases. The predicted structures of both isoforms are shown in Fig. 1.

**Figure 1:**
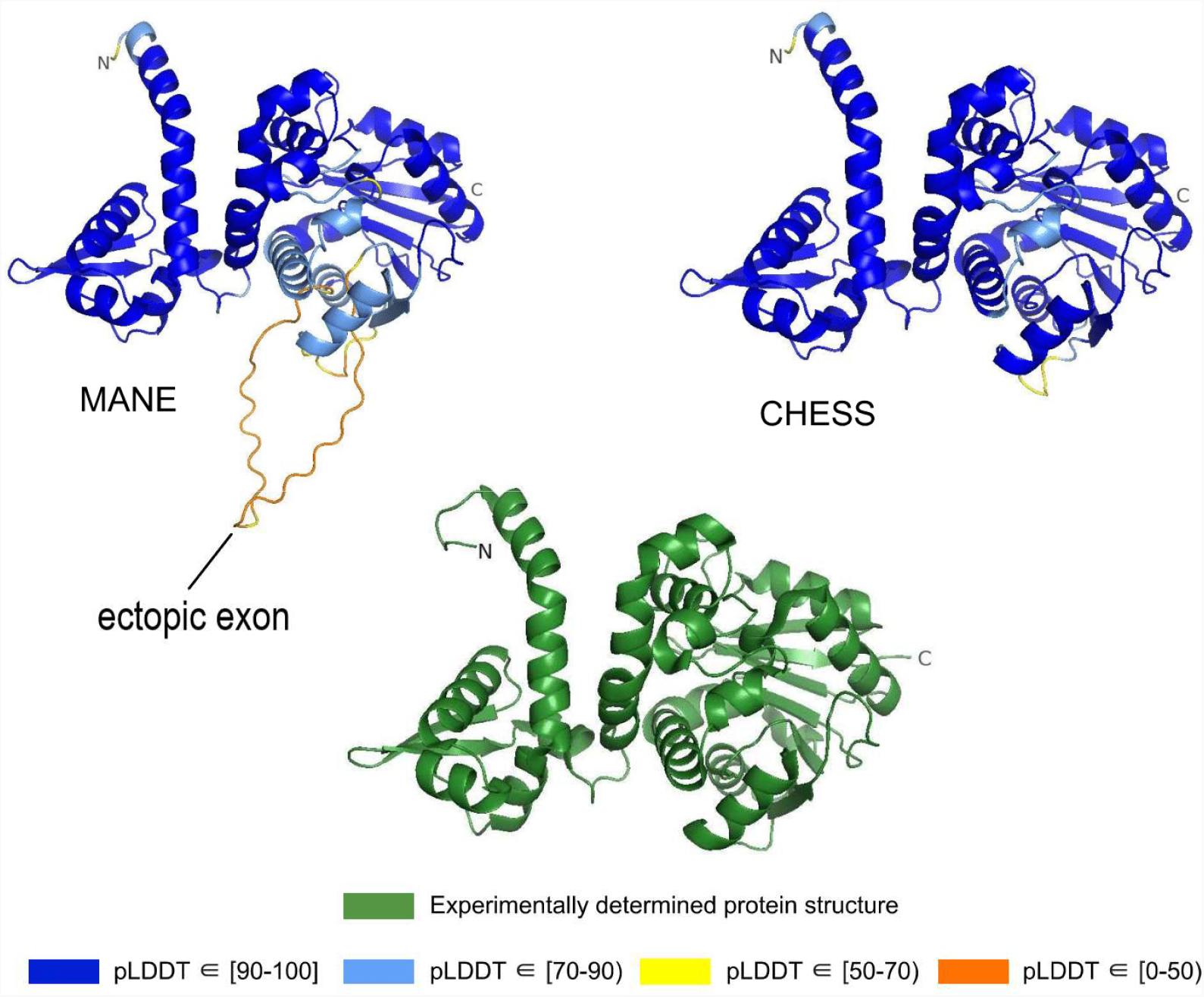
*ASMT* isoform comparison. Comparison of predicted structures of acetylserotonin O-methyltransferase (*ASMT*), showing the 373aa isoform from MANE (CHS.57426.2, RefSeq NM_001171038.2, GENCODE ENST00000381241.9) on the left, and a 345aa alternate isoform from CHESS (CHS.57426.4, GENCODE ENST00000381229.9) on the right. The CHESS 345aa isoform closely matches the experimentally determined X-ray crystal structure of the biologically active protein (*20*), shown at the bottom.

We hypothesized that the highest scoring isoform of *ASMT* according to AlphaFold2 corresponds to the biologically active version of the protein. The score we use for these comparisons is the predicted Local Distance Difference Test (pLDDT) score, which has been demonstrated to be a well-calibrated, consistent measure of protein structure prediction accuracy (*12, 13*). A pLDDT score above 70 (the maximum is 100) indicates that a predicted structure can generally be trusted, while a score below 70 may indicate folding prediction failure or intrinsic disorder within a protein. Scores above 90 imply structure predictions accurate enough for highly shape-sensitive tasks such as chemical binding site characterization. The structure of the 345aa isoform has a very high pLDDT score of 94.7, versus the somewhat lower score of 87.1 for the 373aa MANE isoform.

Because melatonin is primarily synthesized within the human pineal gland at night, we quantified *ASMT* isoform expression using RNA-sequencing data from a previously-published experiment that used tissue extracted from the pineal gland of a patient who died at midnight (*21*). In this tissue sample, the 345aa isoform of *ASMT* was expressed at a level of 327 transcripts per million (TPM), while the 373aa isoform from MANE was expressed at 34 TPM, nearly 10 times lower, supporting our hypothesis that the higher scoring 345aa isoform is functional.

Further evidence for the functionality of the 343aa isoform is shown in Fig. 1. An ectopic exon in the MANE *ASMT* protein creates an unstructured loop that bulges out from the primary structure. The alternate isoform, missing this ectopic exon, closely matches the experimentally determined *ASMT* X-ray crystal structure of the biologically active protein. Furthermore, as reported in Botros et al. (*20*), the insertion of exon 6, corresponding to the ectopic exon in the MANE isoform, distorts the structure and destroys its ability to bind S-adenosyl-L-methionine and to synthesize melatonin. Thus, the structural comparison, the expression evidence, and a melatonin synthesis activity assay all combine to support our hypothesis that the 345aa isoform represents the primary biologically functional isoform of *ASMT*.

#### Gamma-crystallin N

Gamma-crystallin N (*CRYGN*) is a highly-conserved member of the crystallin family of proteins, responsible for the transparency of the lens and cornea in vertebrate eyes (*22*). The intron-exon structure of *CRYGN* has been conserved across at least 400 million years of vertebrate evolution, with close orthologs present in the genomes of chimpanzees, mice, frogs, and the white-rumped snowfinch. Given this extensive evolutionary history, Wistow et al. 2005 (*23*) were surprised to observe that the primate *CRYGN* gene has lost its canonical stop codon, leading them to conclude “the human gene has clearly changed its expression and may indeed be heading for extinction.”

As shown in Fig. 2a, the MANE isoform (CHS.52273.5, RefSeq NM_144727.3, GENCODE ENST00000337323.3) that matches descriptions by Wistow et al. (*23*) includes sequences that do not fold well, as indicated by its pLDDT score of 67.7. However, we found an alternate *CRYGN* isoform, assembled from GTEx (*17*) data, that had a far higher pLDDT score of 92.2, shown in Fig. 2b. Small differences between pLDDT scores may not be meaningful, but large score differences, such as the 24-point gap between the two isoforms of *CRYGN* discussed here, represent a substantial difference in prediction confidence across a large portion of the protein. The higher-scoring *CRYGN* isoform is present in CHESS (CHS.52273.9) and GENCODE (ENST00000644350.1), and it was also present in RefSeq v109 (XM_005249952.4) but was removed in the next release, v110.

**Figure 2:**
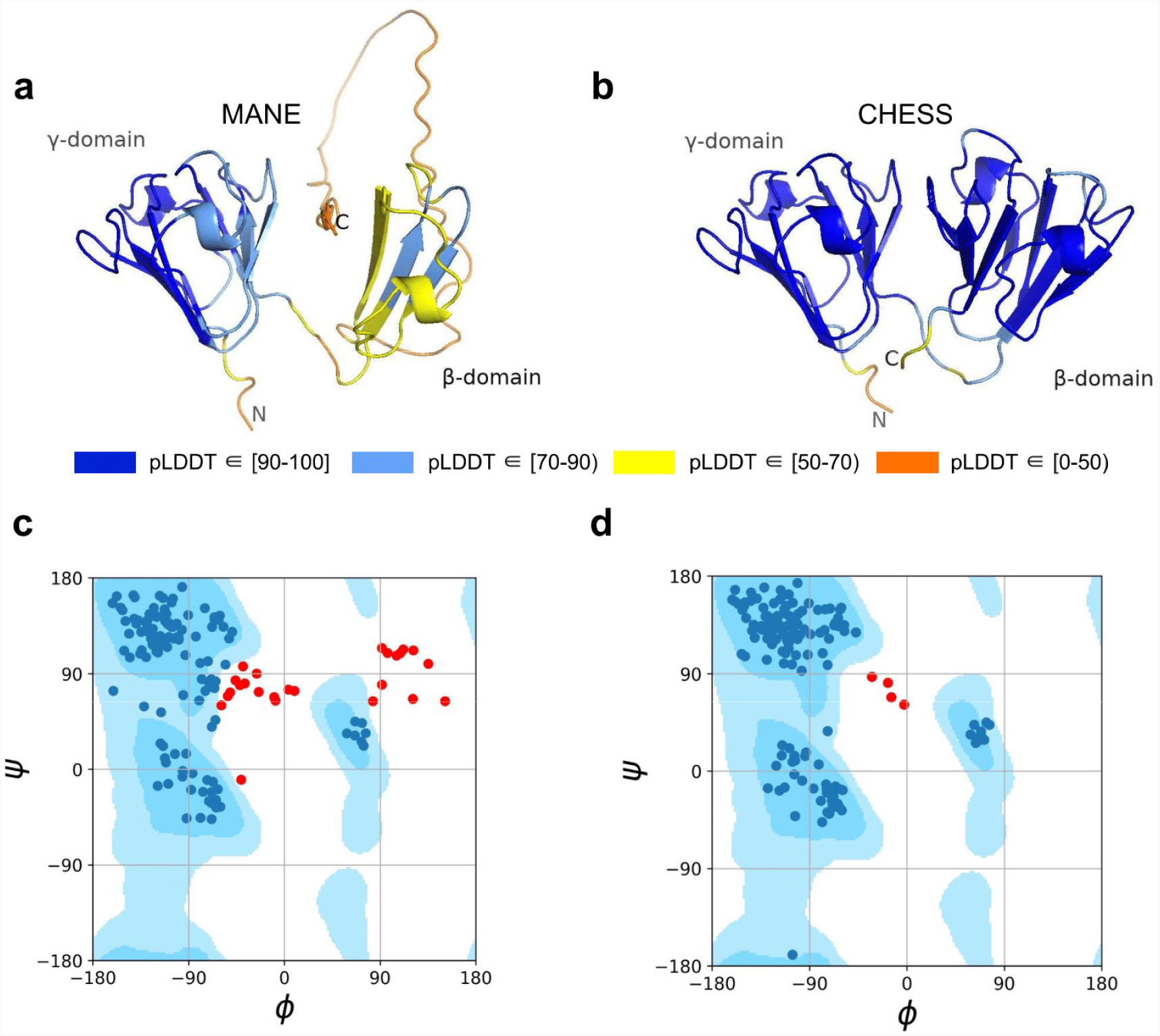
*CRYGN* isoform comparison. (**a**) Predicted protein structure for the MANE isoform (CHS.52273.5, RefSeq NM_144727.3, GENCODE ENST00000337323.3) of Gamma-crystallin N (*CRYGN*), colored by pLDDT score. (**b**) Predicted protein structure for a *CRYGN* alternate isoform (CHS.52273.9, GENCODE ENST00000644350.1). (**c**) Ramachandran plot for the MANE (CRYGN) isoform. Dark blue areas represent “favored” regions while light blue represent “allowed” regions (*24*). The 32 red dots represent amino acid residues with secondary structures that fall outside the allowed regions. (**d**) Ramachandran plot for the alternate *CRYGN* isoform with 4 red dots falling in disallowed regions, compared to 32 disallowed in MANE. All residues associated with the 4 red dots in the alternate isoform are shared with the MANE isoform in the poorly folded N-terminal region.

Both the MANE and alternate isoforms are exactly the same length despite having different C-terminal sequence content. Visual comparison of the predicted structure for the alternate isoform reveals a marked improvement in the structure in the β domain and a clear recovery of *CRYGN*’s dimer-like characteristic, with two structurally similar domains as shown in Fig. 2b. Ramachandran plots (Fig. 2c,d) also support the structure of the alternate CHESS isoform.

Encouraged by the substantially improved folding of this alternate isoform, we examined the intron-exon structure of *CRYGN* in human to determine how it recovered its functional shape despite losing its original stop codon. We found that the common vertebrate four-exon structure of *CRYGN* has changed to a five-exon structure in humans, as shown in Fig. 3. In the well-folded alternate isoform, a novel primate-specific splice site removes the last four amino acids as compared to other vertebrates, but the new primate-specific fifth exon contains a downstream stop codon that adds four residues. The poorly folding MANE isoform (Fig. 3, bottom), in contrast, entirely skips the fourth exon, resulting in a frameshift that adds 43 C-terminal amino acids which have no homology to any *CRYGN* sequence outside of primates.

**Figure 3:**
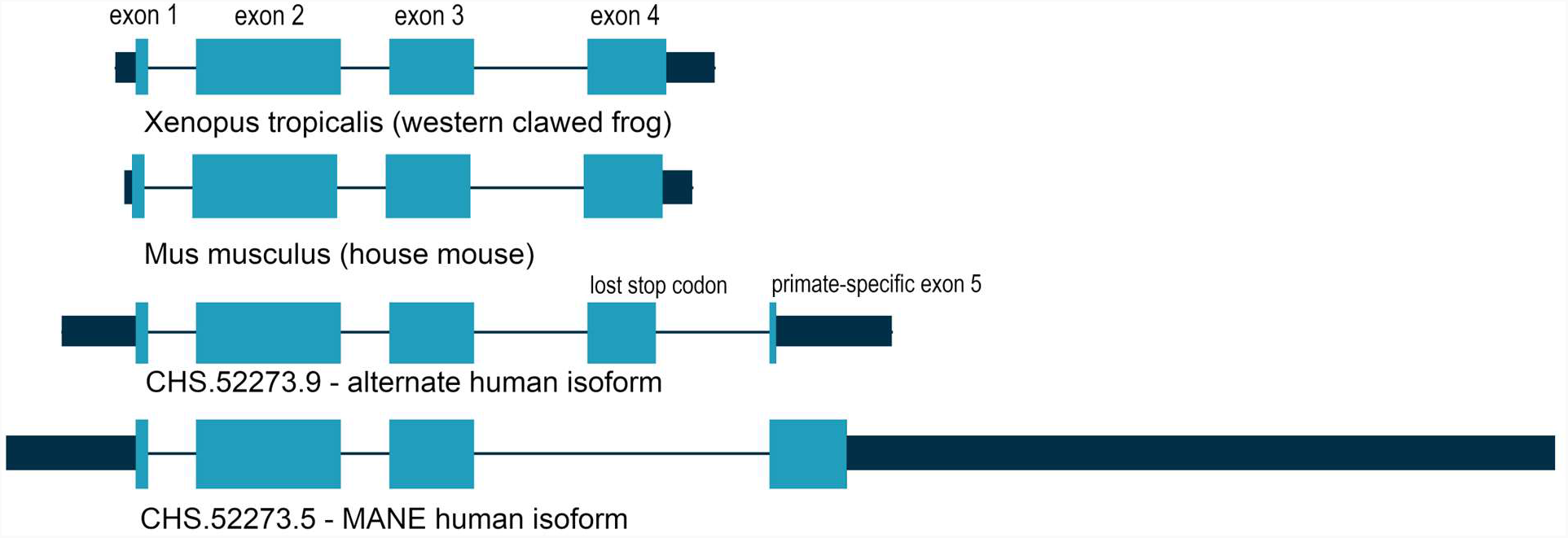
*CRYGN* intron-exon structure. Comparison of Gamma-crystallin N (*CRYGN*) gene structure in frog, mouse, and human. Exons 1, 2, and 3 are highly conserved across all species. Exon 4 is missing from the poorly folding MANE isoform, while exon 5 shows no homology to any species outside of primates. The loss of a stop codon in human exon 4 appears to be balanced by the inclusion of a short novel exon that adds only four amino acids to the final protein. Coding portions of exons are shown with thicker rectangles in teal. Intron lengths are reduced proportionately for the purpose of display.

#### Thioredoxin domain-containing protein 8

The thioredoxin protein family represents an ancient group of highly-conserved small globular proteins found in all forms of life (*25*). Thioredoxin domain-containing protein 8 (*TXNDC8*, alternatively *PTRX3*) is a testis-specific enzyme responsible for catalyzing redox reactions via the oxidation of cysteine from dithiol to disulfide forms (*26*).

As shown in Fig. 4, several canonical protein motifs appear altered or missing in the predicted structure of the human *TXNDC8* MANE transcript, as it lacks a highly-conserved sequence that should start only eight residues away from the CGPC dithiol/disulfide active site. The α2 helix is severely truncated, leading directly to the α3 helix, thereby entirely skipping the β3 sheet. The β5 sheet, normally providing an interaction bridge between the α3 and α4 helices, is similarly missing in its entirety. Finally, the α4 helix is present but rotated 140 degrees relative to its canonical position. These large alterations to the fundamental thioredoxin protein structure result in the MANE transcript receiving a pLDDT score of 56.7.

**Figure 4:**
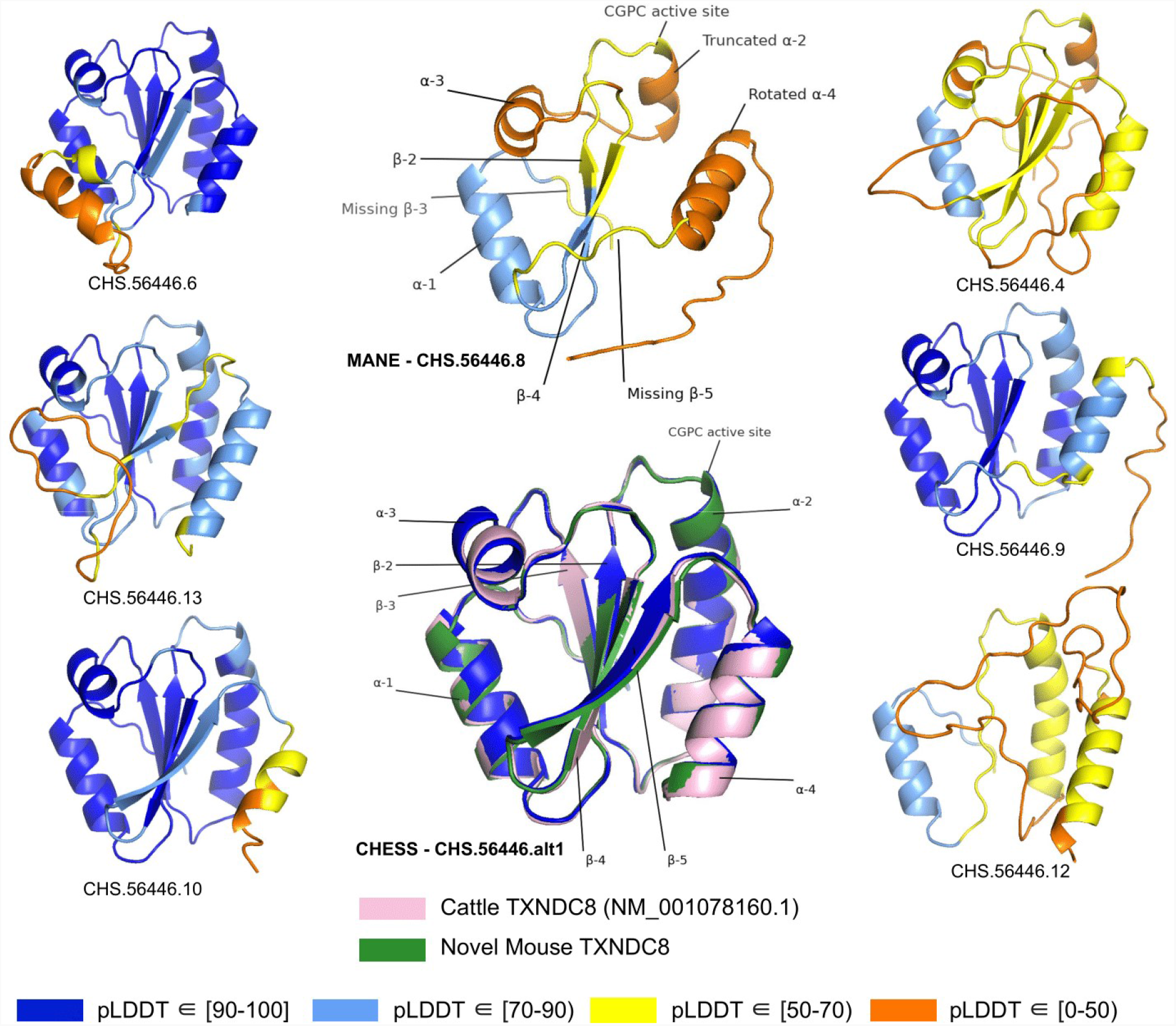
*TXNDC8* isoform comparison. Predicted protein structures for seven distinct human isoforms of thioredoxin domain-containing protein 8 (*TXNDC8*), as well as the primary cattle transcript and a novel mouse transcript. Alternate human isoforms 4, 9, and 12 (right side of figure) lack multiple canonical thioredoxin structures and thus appear non-functional. Several canonical protein motifs are missing or altered in the predicted structure of the MANE transcript (top center). In contrast, the alternate human transcript alt1 matches cattle and mouse to within 0.8 Angstroms. Human transcript CHS.56446.alt1 is colored solid dark blue because every amino-acid residue scores a pLDDT above 90.

In stark contrast to the poorly-folded MANE isoform (CHS.56446.8, RefSeq NM_001286946.2, GENCODE ENST00000423740.7), an alternate isoform assembled as part of the CHESS project (CHS.56446.alt1) and RefSeq (NM_001364963.2) has a pLDDT score of 96.9, an improvement of 40 points. Inspection of the alternate transcript (Fig. 4) reveals full recovery of the canonical α2, α3, and α4 helices as well as the β3 and β5 sheets. Moreover, 3D alignment of the protein encoded by CHS.56446.alt1 to the protein from the primary *TXNDC8* isoform in *Bos taurus* (cow) reveals a very close structural correspondence between the two proteins, with a predicted root-mean-square deviation (RMSD) of 0.8Å. Fig. 4 shows multiple alternative isoforms of human *TXNDC8* from the CHESS annotation, as well as the three-dimensional alignment of CHS.56446.alt1 to its orthologs in cow and mouse. Given its substantially higher pLDDT score and near-perfect structural conservation in other species, the CHESS transcript appears to be a much better candidate for the canonical form of this protein. The MANE isoform, because it is missing multiple key structures, may represent a non-functional product of transcriptional noise.

All isoforms of *TXNDC8* shown in Fig. 4 were assembled from RNA sequencing data during the construction of the CHESS database. This figure illustrates another potential use of structure prediction, namely the ability to distinguish among multiple functional and non-functional isoforms when annotating a genome. As discussed above, the MANE *TXNDC8* isoform appears non-functional, lacking several key structures. In addition, isoform 12 in Fig. 4 appears clearly non-functional, lacking all four of the β sheets and one of the α helices of isoform alt1. Isoforms 4 and 9 also appear likely to be non-functional: both are missing one of the β sheets, and isoform 4 has a low pLDDT score of just 52. Although this example is only one of many, it illustrates how one can employ accurate 3D structure prediction in virtually any species as a powerful new tool to improve gene annotation.

Assemblies of RNA-seq experiments typically reveal thousands of un-annotated gene isoforms, as was illustrated by the use of GTEx to discover more than 100,000 new isoforms when building the original CHESS gene database (*4*). The approach used here, computationally folding each distinct protein encoded by alternate isoforms, allows us to compare the structure of these predicted proteins to the “best” structure for each protein-coding gene locus. For those proteins with at least one high-confidence structure, this strategy may allow us to identify and remove potentially non-functional isoforms.

#### Interleukin 36 Beta

Interleukin 36 Beta (*IL36B*, alternatively *IL1F8, FIL1-ETA*, or *IL1H2*) mediates inflammation as part of a signaling system in epithelial tissue. The pro-inflammatory properties of *IL36B* have been implicated in the pathogenesis of psoriasis (*27*), a common disease characterized by scaly rashes on the skin. *In vitro* and *in vivo* studies have found consistently increased expression of *IL36B* in psoriatic lesions, making the gene a potential target for future anti-psoriatic drugs (*28*).

The ColabFold structure of the *IL36B* MANE isoform (CHS.30565.1, RefSeq NM_014438.5, GENCODE ENST00000259213.9) averaged a pLDDT of only 50.2, a score indicative of near-complete folding failure, while an alternate isoform in CHESS (CHS.30565.4) and RefSeq (XM_011510962.1), shown in Fig. 5, scored 93.0, the largest relative score increase of any isoform we examined. This alternate isoform contains two C-terminal exons that are not present in the MANE isoform and that contribute nearly half of the total protein sequence. A BLASTP homology search of the coding sequence of the final C-terminal exon in the MANE transcript yielded no hits beyond primates. In contrast, a BLASTP search of sequence unique to the alternate isoform revealed significant similarity (e-values of 0.001 and smaller) to *IL36B* orthologs in 1517 organisms including mouse, rat, and Hawaiian monk seal. In addition, expression of the MANE isoform was observed in only 1 sample in the GTEx data with an expression level of just 0.01 TPM, while the CHESS isoform was found in 775 samples with a much higher expression level of 8.4 TPM (Table S3).

**Figure 5:**
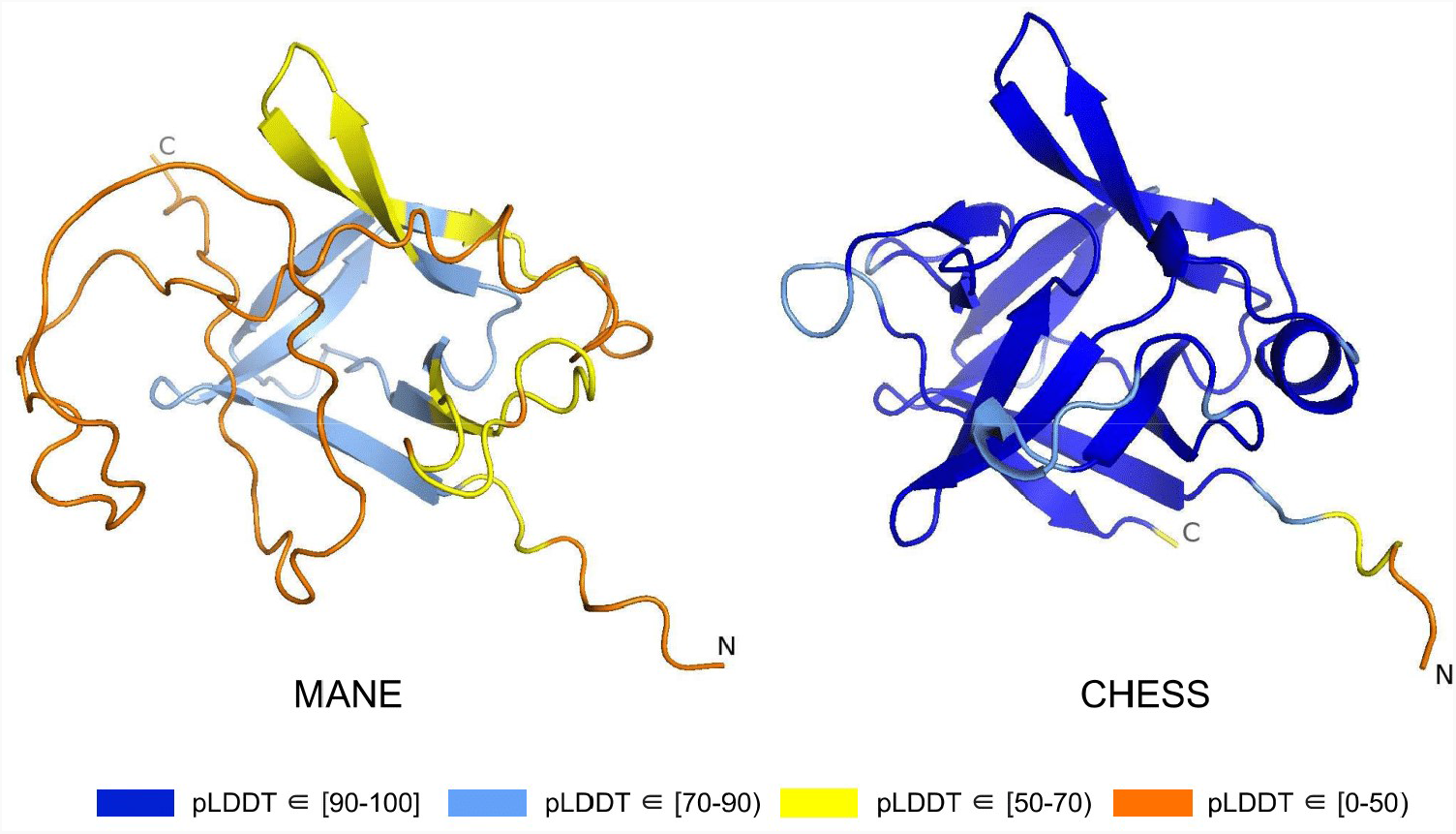
*IL36B* isoform comparison. Comparison of predicted structures for interleukin 36 beta (*IL36B*) for the MANE isoform (CHS.30565.1, RefSeq NM_014438.5, GENCODE ENST00000259213.9) and an alternate isoform from CHESS and RefSeq (CHS.30565.4, RefSeq XM_011510962.1). The highly-conserved protein sequence of the alternate human isoform achieves a very high pLDDT score of 93.0, versus the MANE isoform’s much lower pLDDT of 50.2.

Further probing the functionality of our high-scoring isoform, we aligned the predicted three-dimensional structures for *IL36B* in human, mouse, and rat. As expected, the mouse and rat proteins aligned to each other remarkably well, with a RMSD of 0.60Å. We found the low-scoring MANE protein aligned poorly to the structures for mouse and rat, averaging a distance of 2.74Å, while the alternate isoform aligned far better with an RMSD of 0.76Å. This close similarity in both sequence and structure to conserved orthologs in distant species strongly reinforces the argument that the alternate isoform represents the functional version of the protein in human.

#### Post-GPI attachment to proteins 2

The protein known as Post-GPI Attachment to Proteins 2 (*PGAP2*, alternatively *FRAG1* or *CWH43N*) is required for stable expression of glycosylphosphatidylinositol (GPI)-anchored proteins (*29*) attached to the external cellular plasma membrane via a post-translational modification system ubiquitous in eukaryotes (*30*). Mutations in GPI pathway proteins have been linked to a wide variety of rare genetic disorders (*31*), while mutations in *PGAP2* specifically have been shown to cause to intellectual disability, hyperphosphatasia, and petit mal seizures (*32*).

Out of 85 GTEx-assembled transcripts for *PGAP2* produced during the latest build of the CHESS database, encoding 33 distinct protein isoforms, the single highest scoring isoform according to ColabFold was CHS.7860.59 (RefSeq NM_001256240.2, GENCODE ENST00000463452.6), with a pLDDT of 87.9. The coding sequence of this isoform exactly matches the sequence of the assumed biologically active protein (*32, 33*), and all intron boundaries are conserved in mouse. For comparison, the annotated MANE protein (CHS.7860.58, RefSeq NM_014489.4, GENCODE ENST00000278243.9) has a pLDDT of 78.0 and the intron boundaries are not conserved in mouse. Predicted structures of both proteins are shown in Fig. 6.

**Figure 6:**
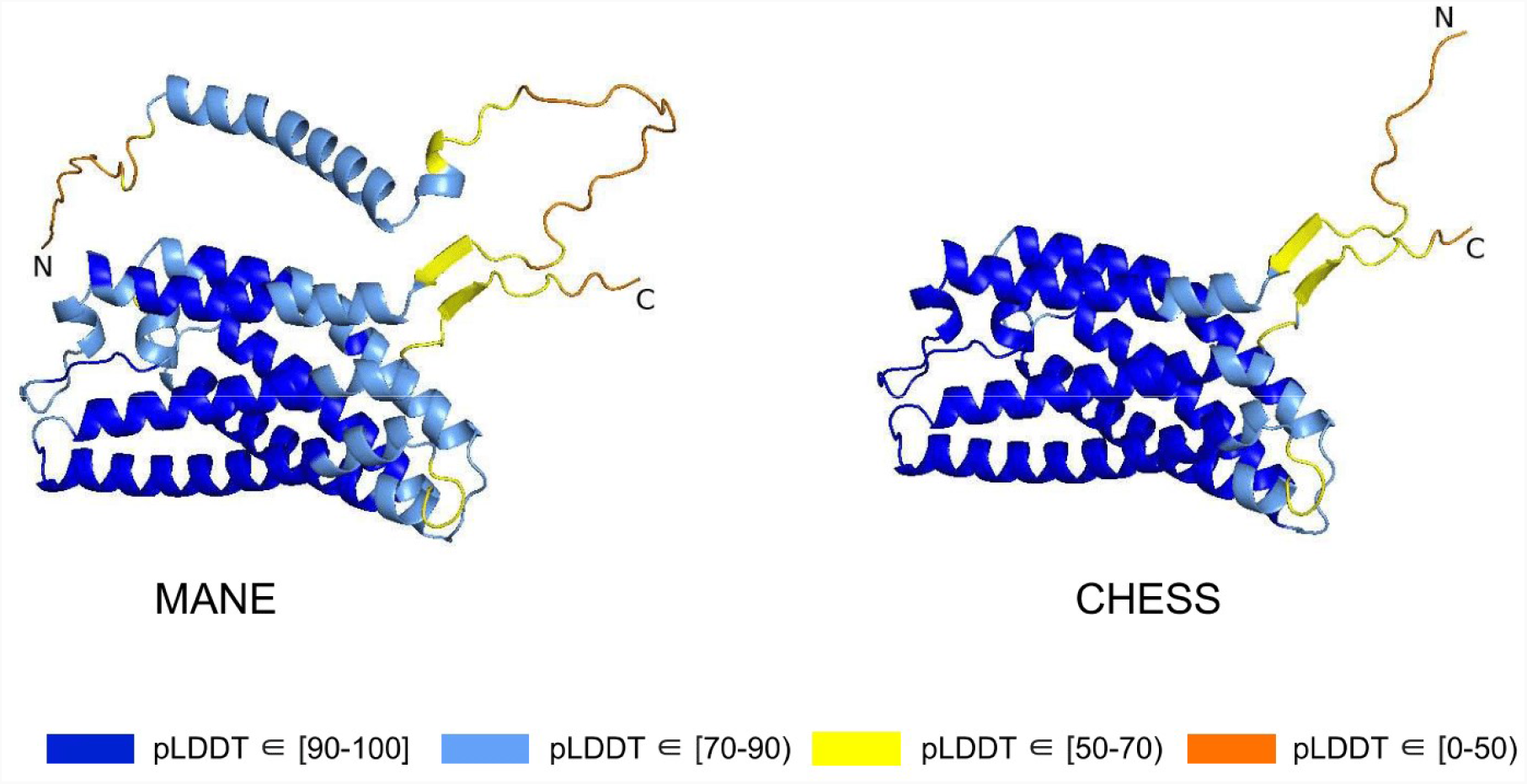
*PGAP2* isoform comparison. Comparison of the structure of the MANE isoform (CHS.7860.58, RefSeq NM_014489.4, GENCODE ENST00000278243.9) versus the highest scoring alternate isoform (CHS.7860.59, RefSeq NM_001256240.2, GENCODE ENST00000463452.6) for PGAP2. Of 33 distinct annotated protein isoforms of *PGAP2*, the one with the highest pLDDT represents the biologically active version (*32, 33*) of *PGAP2* in humans.

RNA-seq data from GTEx also showed that the higher scoring isoform, CHS.7860.59, was expressed in 8776 samples with an average expression level of 2.6 TPM, compared to only 4116 samples with an 0.9 TPM average for the MANE isoform. Comparing their intron-exon structure revealed that the MANE transcript has one extra exon (the second exon out of six). On average across all 31 tissues in the GTEx data, five times more spliced reads supported skipping that exon, as in CHS.7860.59, rather than including it.

### Functional splice variants may not fold well

Alternative splicing allows genes to code for multiple functional protein products (*34*). Thus, rejecting all but the top-scoring isoform based on predicted structure may eliminate lower-scoring yet functional proteins. The risk of discarding functional transcripts by relying too heavily on the pLDDT score is well-illustrated by vascular endothelial growth factor B (*VEGFB*), a growth factor implicated in cancer and diabetes-related heart disease (*35*). The human *VEGFB* gene encodes two well-characterized protein isoforms: *VEGFB-167* and *VEGFB-186*. Alternative splicing that skips part of the sixth exon in *VEGFB-167* leads to sequestration of the protein to the cell surface due to a highly basic C-terminal heparin binding domain. Full inclusion of exon six in *VEGFB-186* results in a soluble protein freely transported to the blood stream (*36*).

Both isoforms shown in Fig. 7 represent highly-expressed and similarly functional products, containing a well conserved cysteine-knot motif (*37*), yet *VEGFB-167* receives a pLDDT score of 81.7 while *VEGFB-186* receives a much lower pLDDT score of 69.5. The MANE isoform (CHS.9039.1, RefSeq NM_003377.5, GENCODE ENST00000309422.7) encodes the freely-soluble protein *VEGFB-186*, while the alternate isoform (CHS.9039.2, RefSeq NM_001243733.2, GENCODE ENST00000426086.3) encodes the sequestered protein *VEGFB-167*. Additionally, *VEGFB-186* is present as a full-length cDNA clone (MGC:10373 IMAGE:4053976) in the Mammalian Gene Collection (*38*), a fact which strongly supports its functionality. Due to the large pLDDT score difference between the two functional *VEGFB* isoforms, a naïve attempt to use protein folding prediction scores as the sole oracle of protein function might inadvertently discard *VEGFB-186*, a clearly functional transcript. Thus, one must be careful to incorporate multiple sources of information when making decisions about isoform functionality.

**Figure 7:**
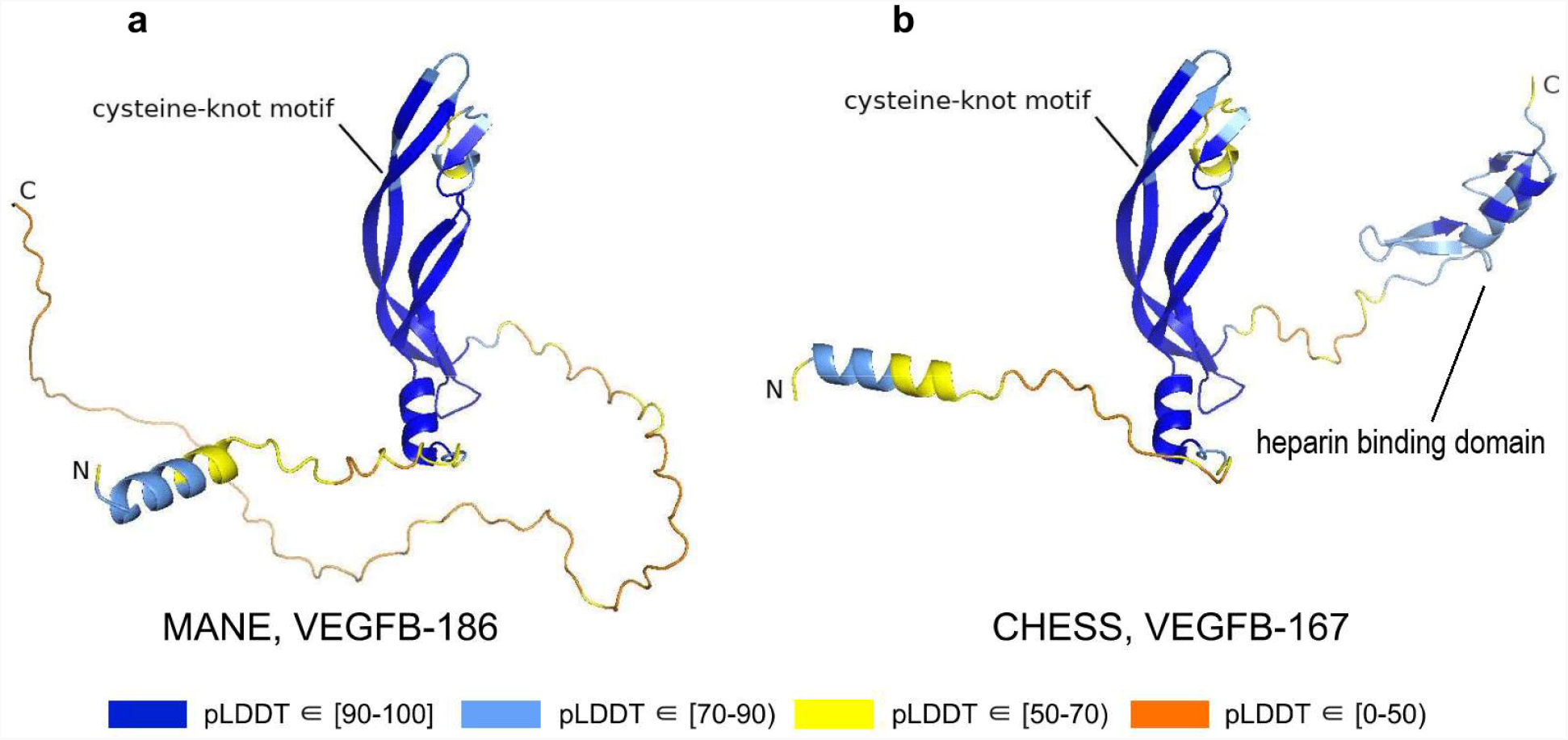
*VEGFB* isoform comparison. Vascular endothelial growth factor B (*VEGFB*) isoforms *VEGFB-186* (**a**) and *VEGFB-167* (**b**). The inclusion of a heparin binding domain in *VEGFB-167* results in sequestration to the cell surface while *VEGFB-186* remains freely soluble. Relying solely on pLDDT comparisons in this case would be misleading, as both isoforms represent well-understood functional protein products.

### A novel protein-coding transcript in mouse

While examining the evolutionary conservation of the three-dimensional structure of *TXNDC8* in human, we noticed that the predicted structure for the same gene in *Mus musculus* (house mouse) seemed to contain a poorly folded region similar to CHS.56446.6, a low-scoring human protein. Further inspection of *TXNDC8* in the mouse genome revealed that the primary RefSeq transcript contains a misfolding fifth exon, while the alternate RefSeq transcript skips the misfolding exon, similar to the functional human isoform, shown in Fig. 8. Interestingly, both mouse transcripts contain a third exon homologous to the sequence missing in the human MANE isoform. A BLASTP search of the misfolding exon resulted in only a single significant hit outside the order *Rodentia* to an unnamed protein product. In contrast, a BLASTP search of the third exon present in all mouse transcripts, homologous to the missing exon in the human MANE transcript, revealed significant hits to at least 31 homologs outside *Rodentia*.

**Figure 8:**
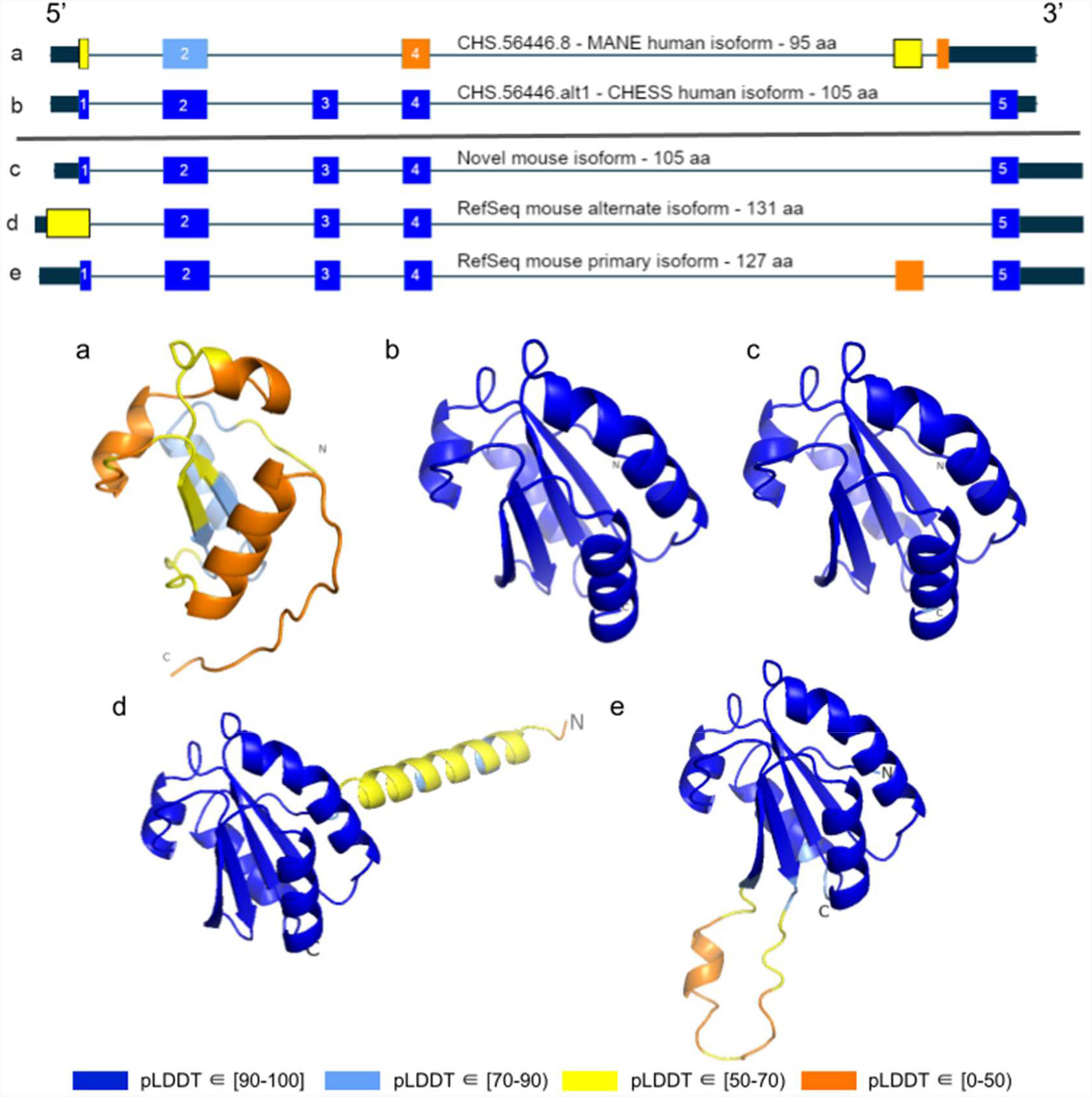
TXNDC8 human and mouse comparison. Intron-exon and predicted protein structure for TXNDC8 in human (**a** and **b**) and mouse (**c, d**, and **e**). Exons are colored according to their average pLDDT score. The highest-scoring isoforms in both human (**b**) and mouse (**c**) share conserved intron-exon structure and nearly identical predicted protein structure.

Unlike the functional human isoform, however, the RefSeq alternate mouse transcript is annotated with a start site 26 codons upstream of the translation initiation site annotated in the human orthologs. A BLASTP search of the 26 amino-acid additional sequence resulted in zero significant hits outside *Rodentia*. Folding this alternate mouse transcript revealed that these additional N-terminal amino acids fail to form any confident predicted protein structure. As a result, none of the three transcripts annotated in mouse fold into the highly-conserved structure of *TXNDC8* in human. Predicted structures and exon alignments for isoforms in human and mouse are shown in Fig. 8.

We hypothesized that a truly functional isoform of *TXNDC8* in mouse should be similar to the human isoform in three-dimensional structure. Based on our observations, this similarity might be realized if the mouse alternate isoform simply started at the downstream start site that matches human, yielding a 105aa protein rather than the 131aa protein that is currently annotated. Folding the coding sequence of the alternate mouse transcript, minus the 26 N-terminal amino-acid residues, revealed a predicted structure remarkably similar to both human and cattle *TXNDC8*. Fig. 4 shows the three-dimensional alignment of the human and cow proteins to our predicted (105aa) isoform of mouse *TXNDC8*. Remarkably, the predicted average RMS deviation between aligned heavy atoms of the putative human and mouse proteins is just 0.83Å. For reference, the atomic diameter of one carbon atom is 1.4Å.

In a further investigation of mouse *TXNDC8* transcription, we aligned 8 gigabases of RNA-seq cDNA from a mouse testis sample (SRR18337982) to the GRCm38 reference genome using HISAT2 (*39*) then assembled transcripts using StringTie2 (*40*). This resulted in two putative transcripts at the mouse *TXNDC8* locus, with neither transcript containing the upstream start site present in the RefSeq annotation. Examination of the read coverage confirmed that both putative *TXNDC8* transcripts appear to use the start site of our proposed shorter protein-coding sequence, with 1060 reads supporting the canonical start site and zero reads supporting the upstream start site. As predicted, one of these newly assembled transcripts contained the protein-coding sequence necessary to exactly match our hypothesized structure-conserved isoform.

We believe this represents the first experimentally confirmed novel isoform in any organism discovered due to a hypothesis derived from comparison of computationally predicted protein structures. All in all, the structure-guided identification and subsequent experimental confirmation of a novel functional *TXNDC8* isoform in mouse demonstrates the power of three-dimensional protein structure prediction to enhance functional annotation in any genome.

### A resource for human annotation

The examples discussed here are only a small subset of the 90,413 structures we generated representing 20,021 human gene loci. We have provided all of these structures as a searchable and downloadable database, at isoform.io, to create a public resource for improving the annotation of the human genome. We provide structure predictions for all isoforms of protein-coding genes shorter than 500aa that we assembled as part of our ongoing effort to improve the CHESS human gene catalog. All CHESS isoforms in this collection have direct support from RNA-sequencing data, as they were assembled from the large GTEx collection, a high-quality set of deep RNA-sequencing experiments across dozens of human tissues. A very small number of MANE proteins were not assembled from GTEx data, but structures of these too are included so that no MANE genes would be omitted.

In the online resource, we provide for each transcript: (1) the nucleotide and amino-acid sequences; (2) the predicted structure, as a file that can be viewed in a standard structure viewer such as PyMOL (*41*); (3) the pLDDT score of that structure; (4) the length of the isoform; (5) the number of GTEx samples in which the isoform was observed; (6) the maximum expression of the isoform in any tissue; (7) an indicator based on alignment of whether all introns are conserved in the mouse genome; (8) an interactive table with functionality to search and sort transcripts to find isoforms of interest; and (9) a Foldseek (*42*) interface to search any given protein structure against all 142,280 transcripts presented here. These predictions can be mined to discover, for example, cases where a known protein gets a surprisingly low pLDDT score, or where alternative isoforms have structures that get higher scores and appear more stable than previously-reported forms of the same protein.

## Discussion

In this analysis, we demonstrated the ability to improve protein-coding gene annotation by predicting three-dimensional protein structures. We searched tens of thousands of predicted structures of alternate isoforms of human genes, and identified a subset that appear to fold more confidently than the isoforms found in MANE, a recently-developed “universal standard” for human gene annotation (*15*). We found hundreds of gene isoforms, all of which were assembled from and therefore supported by RNA sequencing data, that outscored the corresponding MANE transcript.

In the illustratory examples described here, we provide biological and evolutionary context for cases where an alternate human isoform appears clearly superior in structure to its canonical protein. Given the many additional high-scoring transcripts that we identified (Table S3), we expect further improvements in human annotation are yet to be discovered. More generally, we followed a structure-guided annotation strategy that may prove useful in refining the annotation of many non-human species as well.

We expect computational protein structure prediction to become an indispensable tool for future transcriptome annotation efforts. Still, the functionality of many proteins may not be revealed by structure prediction alone. Cases where substantial portions of a protein fail to form a stable structure, such as intrinsically disordered proteins, were not examined here. Although we restricted our analysis to whole-protein comparisons, comparing local structural portions of a protein, potentially near shape-sensitive ligand binding sites (*43*), may enable similar analysis in these proteins. Further advances in predicting structures for multi-chain protein complexes (*44*), as well as improvements in prediction efficiency in large proteins, may expand the range of genes that may be analyzed. An important caveat is that in some cases, truly functional isoforms may receive low predicted folding scores relative to well-folded functional alternate isoforms within the same gene. Thus, structure prediction alone is not always sufficient to make functional claims about any individual protein isoform.

## Methods

### Protein structure prediction

We folded all transcripts in the Comprehensive Human Expressed SequenceS (CHESS) annotation less than 500 amino acids in length. Similar to the initial effort to fold the human proteome (*13*), the length limit was chosen to make the overall computational runtime feasible. This yielded 142,280 transcripts representing 90,413 unique protein-coding sequences at 20,021 loci. Coding sequences for CHESS transcripts were determined with ORFanage (*45*) (commit 7337f4d). For each protein sequence, we generated a multiple sequence alignment by aligning them with ColabFold’s MMseqs2 (*46*) workflow (colabfold_search) against the UniRef100 (*47*) (2021/03) and ColabFoldDB (2021/08) database. Structure predictions were made with ColabFold (commit 3398d3) using AlphaFold2 and MMseqs2 version 13.45111. To speed up the search, we set the sensitivity setting to 7 (-s 7). We predicted each structure using colabfold_batch and stopped the process early if a pLDDT of at least 85 was reached by any model (--stop-at-score 85) or if a model produced a pLDDT less than 70 (--stop-at-score-below 70). All models were ranked by pLDDT in descending order. Run-time was estimated from a sample of 500 proteins randomly selected from the 90,413 structures. Prediction of all structures on 8 x A5000 GPUs required 34 days. Multiple sequence alignment took 34 hours using MMseqs2 on an AMD EPYC 7742 CPU with 64 cores.

### Filtering MANE comparisons

To generate the 401 protein isoforms in Table S3, we used the following filtering criteria and procedures. For each isoform with a distinct coding sequence located at a MANE v1.0 locus and with coding sequence overlapping a MANE annotated protein, we compared the pLDDT score to that of the associated MANE protein. As described previously, pLDDT is a reliable measure of the confidence in a structure, where predictions with 70 ≤ pLDDT ≤ 90 are confident, those with pLDDT > 90 are highly confident, and those below 50 represent low-confidence structures and may be disordered proteins (*48*). We only considered alternative isoforms that had a pLDDT score ≥ 70, indicating a generally well-folded protein, to avoid including any intrinsically disordered proteins (*49*). Filtering and general analysis was performed in Colab (https://colab.research.google.com) with Python version 3.7.13.

We selected isoforms that, when compared to the MANE isoform for the same gene, scored at least 5% higher as measured by pLDDT and were at least 90% as long. Additionally, to capture cases where the MANE transcript might be missing functional sequence elements, we selected alternate isoforms that were at least 5% longer than the MANE isoform, that had equal or higher pLDDT scores, and that were assembled in an equal or higher number of GTEx samples. Finally, to capture cases where an alternate isoform might be functional despite being substantially shorter than the MANE protein, we selected cases with no length threshold where the alternate isoform scored at least 5% higher and was assembled in an equal or higher number of GTEx samples. After applying these filters, we observed that in some cases, a processed pseudogene (*50*) contained within an intron outscored the associated primary transcript. To eliminate such cases, we used GFFcompare (*51*) to ensure that isoforms overlapped their MANE transcript’s coding sequence. When multiple alternate isoforms contained the same coding sequence, and thus received the same pLDDT score, we selected the isoform assembled in the highest number of GTEx samples.

### Annotation sources

Transcript annotations for MANE, CHESS, RefSeq, and GENCODE were retrieved from the following sources. The MANE v1.0 database was downloaded from NCBI at https://www.ncbi.nlm.nih.gov/refseq/MANE. Annotations from the CHESS v2.2 database were retrieved from http://ccb.jhu.edu/chess. Additional CHESS annotations came from an unpublished set of transcript assemblies created as part of the process of building the next major release of CHESS; these were assembled from approximately 10,000 GTEx RNA-seq experiments across 31 tissues using StringTie2 (*40*). Transcripts from this set were given a CHESS ID starting in “hypothetical” if the locus was missing from CHESS 2.2, or else given an ID ending in “altN” if the locus was present in CHESS 2.2 but the exact isoform was not. Note that many of these, particularly those with a poor protein folding score, will not make it into the published CHESS database. RefSeq annotations (releases 109 and 110) were downloaded from https://www.ncbi.nlm.nih.gov/projects/genome/guide/human/index.shtml. GENCODE (v38, v39, v40) annotations were collected from https://www.gencodegenes.org/human/.

### Visualization and atomic alignment

All visualizations and three-dimensional protein structure atomic alignments were performed in PyMol (*41*) version 2.5.2 using non-orthoscopic view, white background, and ray trace 1200,1200. Root-mean-square deviations were calculated without excluding any outliers.

Ramachandran plots were created using PyRAMA version 2.0.2 with Richardson (*24*) standard psi and phi values for all amino acids excluding glycine and proline. Intron-exon structure plots were produced with MISO (*52*) (commit b714021) and TieBrush (*45*) (commit e986d64).

### RNA-sequencing quantification of the human ASMT gene

RNA-sequencing data were downloaded from NCBI for run SRR5756467 from BioSample SAMN07278516, a pineal gland from a patient who died at midnight. A detailed summary of the experimental protocol used to generate these data can be found in NCBI BioProject PRJNA391921. Isoform-level quantification was performed using Salmon (*53*) version 1.8.0.

### RNA-sequencing assembly of mouse TXNDC8

RNA-sequencing data were downloaded for run SRR18337982 from BioSample SAMN26725167, a tissue sample from the testis of a control mouse. A detailed summary of the experimental protocol used to generate these data can be found in NCBI BioProject PRJNA816862. cDNA reads were aligned to the mm39 reference genome using HISAT2 (*39*) version 2.1.0 then assembled into transcripts using StringTie2 (*40*) version 2.2.1.

### Intron conservation in human and mouse

We assessed the conservation of GT-AG intron boundaries between CHESS human transcripts and transcripts from the GRCm38 mouse reference genome. Data for the human-mouse alignment was extracted from a 30-species alignment anchored on GRCh38 that was downloaded from the UCSC genome browser (*54*). We used MafIO in BioPython (*55*) version 1.71 to check if all intron boundaries were conserved between mouse and human transcripts. In the supplementary tables, a value of 1 in the “introns in mouse” column indicates that all boundaries were conserved, while “0” means that at least one splice site (either a GT at a donor site or an AG at an acceptor site) was not conserved in the alignment.

## Acknowledgments

The authors would like to thank all members of the Salzberg, Pertea, and Steinegger labs, as well as David Lipman for helpful feedback during project conceptualization, Snickers and Queso for providing moral support, and Ben Langmead for publicly hosting bulk data files.

## Funding

National Institutes of Health grant R01-HG006677 (SLS)

National Institutes of Health grant R35-GM130151 (SLS)

National Institutes of Health grant R01-MH123567 (SLS)

National Science Foundation grant DBI-1759518 (MP)

National Research Foundation of Korea grant 2019R1-A6A1-A10073437 (MS)

National Research Foundation of Korea grant 2020M3-A9G7-103933 (MS)

National Research Foundation of Korea grant 2021-R1C1-C102065 (MS)

National Research Foundation of Korea grant 2021-M3A9-I4021220 (MS)

The Creative-Pioneering Researchers Program through Seoul National University (MS).

The funders had no role in study design, data collection and interpretation, or the decision to submit the work for publication.

## Author Contributions

Conceptualization: SLS, MS, MJS

Data curation: MJS, AV, SC, SP, IM

Formal analysis: MJS, SLS

Funding acquisition: SLS, MP, MS

Investigation: MJS, SLS, MP, SC, SP

Methodology: MJS, SLS, MP, MS

Project administration: MJS, SLS

Validation: MJS, SLS, MP

Visualization: MJS, NR

Resources: SLS, MS

Software: MJS, MS, SC, SP

Supervision: SLS, MS

Writing – original draft: MJS, SLS

Writing – review & editing: MJS, SLS, MP, MS

## Data and materials availability

Gene identifiers for all predicted protein isoforms as well as pLDDT scores and evolutionary conservation data from mouse can be found in Table S1. Predicted scores and GTEx expression data for all isoforms overlapping a MANE locus can be found in Table S2. Data for the 401 alternate isoforms with evidence of relatively superior structure, and possibly superior function, can be found in Table S3. Additionally, all data can be downloaded from the project website, isoform.io.

## Supplementary table captions

**Table S1: All isoform summary**. Folding scores from ColabFold for each transcript from a preliminary new build of the CHESS database that contained a protein-coding sequence (CDS) that was under 500aa in length. For transcripts already contained in the published CHESS 2.2 database, the identifier from that database is provided. If the transcript maps to a known gene locus X but is a novel isoform, it is shown with the identifier CHS.X.altY. If a transcript occurs at a novel locus X, the identifier is hypothetical.X.Y, where Y identifies the isoform number. Additional columns show the gene name, the RefSeq ID (release 110), the GENCODE ID (release 40), the pLDDT (folding) score, and a flag indicating whether all intron boundaries (for multi-exon genes) are conserved in the mouse genome.

**Table S2: MANE comparison summary**. Folding scores and additional data for all CHESS transcripts that match genes in the MANE v1.0 dataset, limited to protein sequences under 500aa in length. Transcripts must overlap the annotated CDS of the MANE transcript to be included. Columns include: *CHESS_ID_isoform*, the CHESS identifier of the alternate isoform transcript; *CHESS_ID_MANE*, the CHESS identifier of the MANE transcript at the same locus; *gene*, the gene name; *aa_length_isoform*, the amino-acid length of the alternate isoform’s CDS; *aa_length_MANE*, the amino-acid length of the MANE transcript’s CDS; *length_ratio*, the ratio of the alternate isoform length to the MANE isoform length; *pLDDT_isoform*, the predicted folding score of the alternate isoform; *pLDDT_MANE*, the predicted folding score of the MANE isoform; *pLDDT_ratio*, the ratio of the alternate isoform folding score to the MANE isoform folding score; *GTEx_samples_observed_isoform*, the total number of GTEx samples where the alternate isoform was observed at least once; *GTEx_samples_observed_MANE*, the total number of GTEx samples where the MANE isoform was observed at least once; *GTEx_top_tissue_name_isoform*, the name of the tissue in which the alternate isoform was observed in the highest number of samples; *GTEx_top_tissue_name_MANE*, the name of the tissue in which the MANE isoform was observed in the highest number of samples; *GTEx_top_tissue_TPM_isoform*, the average TPM of the alternate isoform in the named tissue; *GTEx_top_tissue_TPM_MANE*, the average TPM of the MANE isoform in the named tissue; *introns_conserved_in_mouse_isoform*, an indicator of whether introns are conserved between the alternate human isoform and any annotated isoform in the GRCm38 mouse reference genome; *introns_conserved_in_mouse_MANE*, an indicator of whether introns are conserved between the MANE human isoform and any annotated isoform in the GRCm38 mouse reference genome.

**Table S3: MANE comparison summary, filtered subset**. A filtered set of CHESS transcripts compared to MANE according to the criteria detailed in the “Filtering MANE comparisons” section of the Methods. Uses the same column names as Table S2.

